# Creating, curating and querying computer models of biological pathways for Inherited Metabolic Disorders

**DOI:** 10.1101/2023.05.24.542073

**Authors:** Denise N. Slenter, Egon L. Willighagen

## Abstract

Collecting knowledge about Inherited Metabolic Disorders (IMDs) has the potential to support early diagnosis as well as foster research into treatment options. Biological pathway databases can be used to create an overview of relevant articles and books describing IMDs and collect, curate, and summarize knowledge about these disorders. Reuse of the information captured in these databases in research requires the knowledge to be accurate but also machine-readable. WikiPathways is a community-driven project to establish a machinereadable knowledge base of biological processes. We here describe how pathways models from WikiPathways were used to represent the underlying biological mechanisms of many IMDs collected by domain experts over a period of six years. This paper describes a standardized approach to depict IMDs in WikiPathways, shows the current limitations in creating machine-readable disease information, and introduces an approach to support data curation based on these machine-readable biological pathways. Furthermore, several SPARQL-queries were developed to analyze the biological content created in these models. Using this approach, 47 pathways were collected about 345 diseases, involving 877 metabolites, 421 annotated metabolic interactions, 262 genes, and 587 proteins related to these disorders.

## 1.1 Introduction

The critical involvement of most enzymes in various metabolic processes (e.g. synthesis, degradation, and molecular transport) [1] comes with the consequence that a malfunction in any one of these enzymes (or even small enzymatic disturbances) can result in serious consequences. An excess of specific metabolites can be toxic to cells and organs, whereas a lack thereof can lead to growth disturbances thereof. Furthermore, significant changes in metabolite levels can (in)activate downstream pathways [2]. People afflicted by these types of enzyme dysfunctions can be classified as suffering from Inherited Metabolic Disorders (IMDs) or Inborn Errors of Metabolism (IEM) [3]. Diagnosis of patients with IMDs can be performed through one of four methods (differing slightly per country):

1. Prenatal (before pregnancy): prenatal screening for carriers of known disorders [4], which is conducted in specific subpopulations known to share a common ancestry. In The Netherlands, these tests are used for inhabitants of a genetically isolated part of the Netherlands [5], and people from the Ashkenazi Jewish population [6]. Recent advanced in massive parallel sequencing techniques such as Next Generation Sequencing (NGS) [7] are often used as a prenatal screening technique.
2. Antenatal (during pregnancy): Non Invasive Prenatal Test (NIPT) [8] and ultrasound [4] methods, where the first is useful in detecting checking trisomy disorders (affecting chromosomes 13, 18, 21) and known genetic variants, while the latter can be used to visualize abnormal growth of organs.
3. Postnatal (close after birth: neonates): through existing screening programs: neonatal heel prick screening [4], based on altered metabolites found through Mass Spectrometry (MS) measurements on dried blood samples. In The Netherlands, 26 disorders are checked for, out of which 20 are IMDs, whereas in the USA 34 diseases can be diagnosed, out of which 25 are IMDs [4]. The Dutch screening program has recently (1st of June 2022) been updated with a test for spinal muscular atrophy (SMA), and other disorders are under review to be added to the panel.
4. Postnatal (after birth: infancy, childhood, and puberty): Patients suffering from IMDs are admitted to the hospital with (severe) symptoms, often at relatively young ages, which initiates the diagnostic process, where a combination of targeted metabolic testing, clinical setting, and family history are used [9]. The metabolic screening in well-equipped laboratories uses tandem mass spectrometry techniques [10], which can be composed of targeted assays for known disorders or an untargeted method for unknown metabolic profiles [11].

Even though genetic variants testing methods, such as NGS or Whole Exome Sequencing (WES) are used for diagnosis, metabolic measurements are considered more sensitive and specific [12]. A correct and timely diagnosis is needed to start treatment, however, most IMDs are currently not treatable [13]. A potential method to understand IMDs better in order to develop suitable diagnostic methods, as well as finding potential treatment targets, might lie with combining information from genomic, transcriptomic, proteomic, metabolomic, and fluxomic data through pathway and network analysis [14–16]. This study shows how biological pathway drawings of IMDs have been converted to machine-readable computer models, how the curation of these models can be aided with automated tests, and how the existing models can be queried for downstream analysis.

## 1.2 Methods

Open Science approaches [17] were used for the full project as well as the FAIR principles [18], to maximize the interoperability and reusability of these models.

### 1.2.1 Pathway figures and their biology

The backbone of the pathway models was based on the Figures drawn in the 2014 (4th) edition of the clinical genetics reference book *Physician’s guide to the diagnosis, treatment, and follow-up of inherited metabolic diseases* [19]. Each chapter describes a group of IMDs related to a set of metabolic reactions. For each pathway figure, the accompanying overview table of disorders and their link to affected genes were used to construct a model, connecting the metabolic substrate, product, and enzymes involved with the disorder for each enzyme. Additional literature was retrieved for enzymes or metabolites which could not be linked to database entries based on the information provided in the chapter, as well as publications and databases on the disorders.

### 1.2.2 Shaping the data models

Machine-readable versions of several metabolic pathways were created using the pathway editor and curation tool PathVisio (version 3.3.0) [20], storing all pathway model knowledge in the Graphical Pathway Markup Language (GPML, version 2013a) [21], which is based on the XML file format [22]. Biological entities were captured as DataNodes (Figure 1.1a A), which contained at least a textual label and biological type, and were extended with literature references and a unique database identifier (ID) (Figure 1.1a B) if available. Additional information which could not be captured within these settings was stored using free text comments (Figure 1.1a B).

**Figure 1.1:**
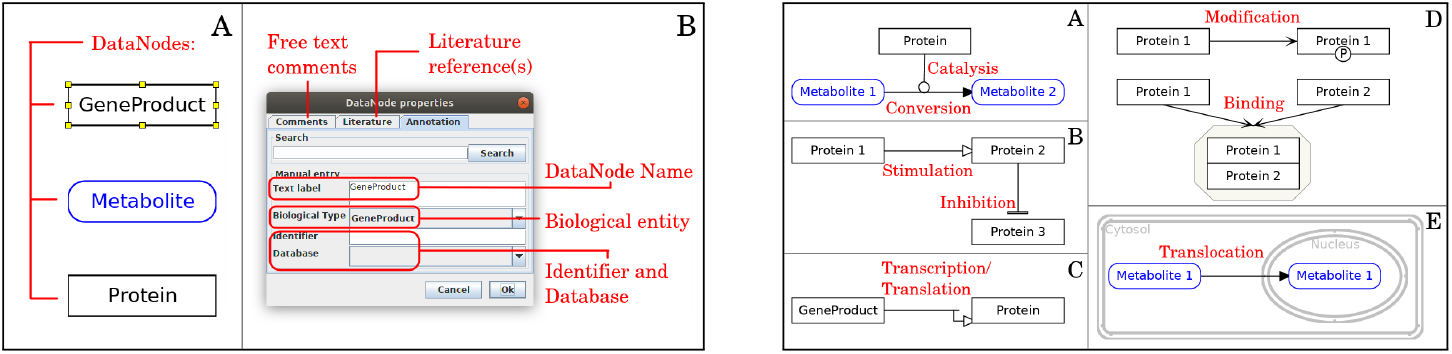
(a) Left Panel: Visualisation of modeling properties for biological entities in PathVisio. A: Main relevant DataNodes, where the GeneProduct is selected (indicated with small yellow blocks). B: Knowledge captured for each individual DataNode, including a database ID, literature reference, and free text comment(s). (b) Right Panel: Overview of MIM-interaction types and how these were used to connect biological entities. Both Figures were adapted from [23].

Interactions between these biological entities were captured with the Molecular Interaction Maps (MIM) standard [24], and if no suitable MIM-interaction was available, with the basic interactions panel available in PathVisio. MIM was used to describe conversion and catalysis (Figure 1.1b A), stimulation and inhibition (Fig. 1.1b B); transcription/translation from gene to protein (Fig. 1.1b C), modification for post-translational or other modifications and binding for complex formation (Fig. 1.1b D); and translocation for transport of metabolites between different cellular compartments (Fig. 1.1b E).

Substrate and product metabolite DataNodes were connected through a MIM-conversion interaction, and connected to their acting enzymes (as a protein DataNode) using an anchor and MIM-catalysis interaction. The same anchor was used to connect a textLabel through a basic interaction arrow, to connect IMDs to the pathways models. Groups of proteins catalyzing the same reaction (without influencing each other) were grouped and connected with one MIM-catalysis interaction, whereas complexes were added by grouping the relevant DataNodes (including relevant co-factors) as a complex. Proteins were annotated through BridgeDb [25], by searching the name of the gene or protein mentioned in the book chapter. The UniProt database [26] was used to retrieve IDs for the metabolic conversions from the Rhea database [27], if available. The metabolite DataNodes were annotated with ChEBI IDs [28] (if available) in line with the entities described in Rhea. The IMD textLabels were annotated using the href option in GPML2013a, by adding IDs from the OMIM database [29]. When these searches did not lead to any results, we explored various databases and online resources, to retrieve suitable IDs. When no relevant ID was found, free text comments were used to describe the entity.

A visual legend was added for each individual pathway, describing the DataNodes and interaction types used for that pathway. The entities within the legend were added as graphical elements (to avoid issues with downstream analysis). A textual description of the pathway was added, describing the content of the whole pathway, including a reference to the relevant book chapter [19] and additional noteworthy details of the disorders described within the model.

### 1.2.3 Disseminating the pathway models

The pathway models were uploaded to WikiPathways [30], which provides them with a unique ID (WPxxxx) and URL for online visualization of the pathway. Relevant Pathway Ontology [31] and Human Disease Ontology [32] terms were added through the WikiPathways website (wikipathways.org), using an ontology tagging system sourced from BioPortal [33]. The models were annotated with Quality Assurance tags (e.g. ‘Approved for Data Analysis’), and relevant community tags (e.g. ‘Inborn Errors of Metabolism (IEM)’). The community portal (imd.wikipathways.org) was updated with new pathways, including their status (‘Approved’ or ‘In Progress’), and information on their chapter numbers for both edition 4 and edition 5 of the book [34].

Through the Pathway Widget [35] interactive pathway visualizations of the IMD models (based on their community tag) are depicted on the IEMBase database [36], highlighting the protein of interest for each individual disorder, as well as other pathways containing this protein (or respective gene). The visualizations are created from the Scalable Vector Graphics (SVG) format (github.com/wikipathways/tool pathway - viewer), and use syntax highlighting of the relevant protein based on the label of the DataNode (classic.wikipathways.org/index.php/PathwayWidget).

### 1.2.4 Semantic web conversion of pathway models

To support computer-assisted data curation, the pathway models were converted to the Resource Description Framework (RDF) format [37, 38] to harmonize the data from all models (github.com/wikipathways/GPML2RDF). This RDF consists of two parts: a conversion of the native GPML format to RDF, capturing all ‘raw’ data in a semantic manner (called GPML-RDF); and the harmonized version of the former, which reduces the RDF to a unified model and enhances the data with additional information such as ID mapping through BridgeDb [25] (called WP-RDF). Figure 1.2 exemplifies one interaction connecting a disorder to a metabolic conversion, and how this information is stored in GPML-RDF and WP-RDF. The IMDs captured as textLabels are currently only available in the GPML-RDF and can be connected to the WP-RDF content through either the Rhea ID or the unique interaction reference that is assigned by PathVisio.

**Figure 1.2:**
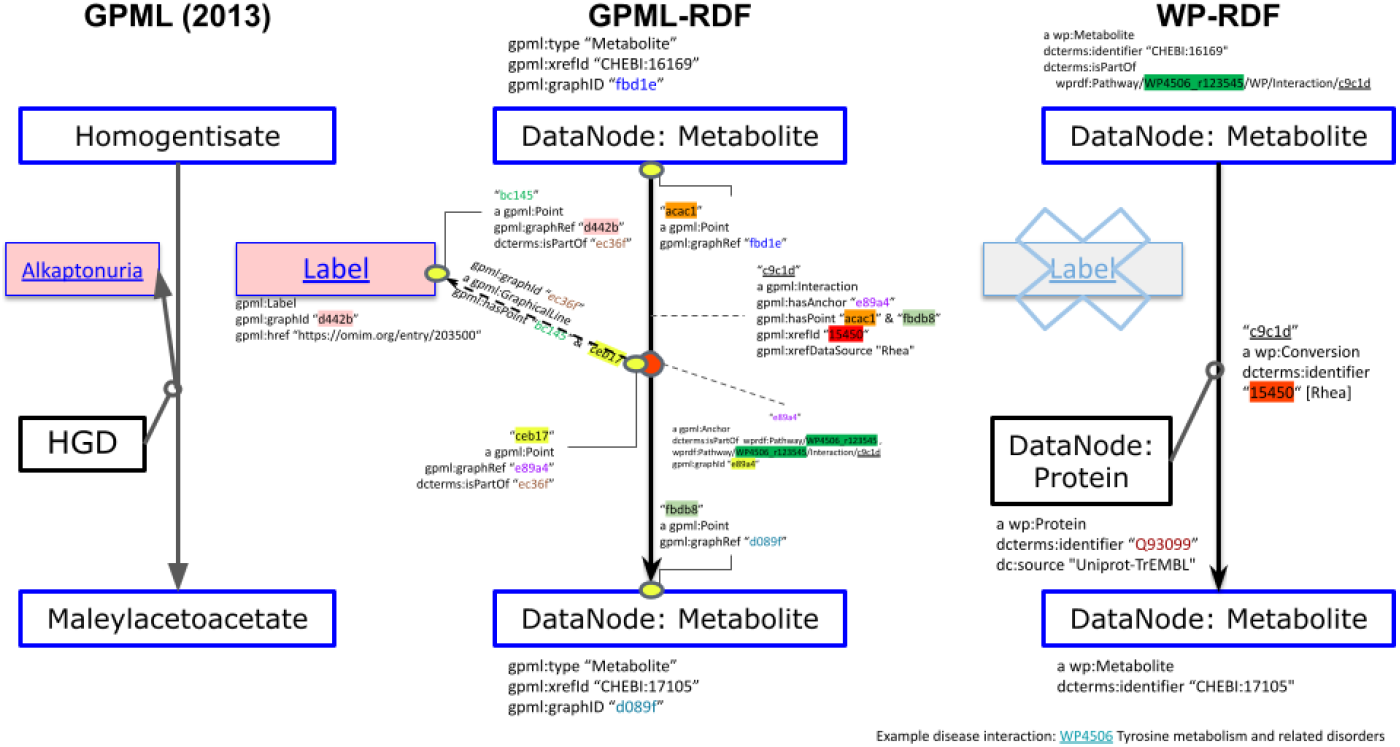
Pathway model data captured in GPML(2013), GPML-RDF, and WP-RDF schema, in the case of one substrate (homogentisate) being converted to one product (maleyl acetoacetate) by one protein (HGD), which is linked to the IMD alkaptonuria.

Originally, not all information relevant to the IMD pathways was harmonized in the WP-RDF. Therefore, several updates were accomplished to the original GPML2RDF code: adding the community tag (‘Curation:IEM’) to allow filtering of pathway model content, and the addition of RHEA IDs to the RDF output. Furthermore, to support the integration of metabolomics and chemical biomarker data, neutral InChIKeys [39] were added to the WP-RDF.

### 1.2.5 Automated testing the content

The generated RDF for the biological pathways was used to support the curation using an automated curation process. Starting in 2013, a Java library was developed [23] where a combination of SPARQL and JUnit tests were used to test the content of the pathway models (github.com/wikipathways/WikiPathwaysCurator).

Tests were implemented on a continuous basis (see Table 1.1). Several tests were written to detect problems directly in the RDF through SPARQL queries (list the disease labels in the pathways found in the source code repository, see Code Example 1), other to check problems in the content returned by the SPARQL query. Two newly developed test relevant to IMD pathway models checks the annotations of diseases in the pathways with links to the OMIM database. The first test checks that an external link is given for the disease label, while a second checks the format of the link if made to OMIM, see Code Example 2.

**Table 1.1:** Overview of automated tests, categorized by Type and Theme, with examples of specific tests. The COVID-19 and LIPID MAPS tests are specific for those communities and implement FAIR maturity indicators defined by these communities (R1.3).

| Type | Theme | Count | Example |
| --- | --- | --- | --- |
| GeneProduct identifiers | Ensembl identifiers | 5 | Test for old Ensembl identifiers, and for species mismatches. |
|  | UniProt | 5 | Test for unreviewed proteins and deleted identifiers. |
|  | Other | 5 | Test for ID format mismatches. |
| Metabolite identifiers | CAS registry numbers | 2 | A test for deleted CAS registry. |
|  | ChEBI IDs | 4 | A test for replaced ChEBI identifiers. |
|  | HMDB IDs | 5 | A test for the correct format. |
|  | Other | 24 | ChemSpider IDs must be integers. Metabolites must not have gene identifiers. |
| Interactions |  | 8 | Tests that proteins are not converted into metabolites (except hormones). |
| Pathway |  | 19 | Tests for ontology annotations and title format. |
| References |  | 4 | Cites a retracted article. |
| Other | identifiers | 5 | Tests that check any BridgeDb mappings are made. |
|  | DataNodes | 4 | A test that all DataNodes have an identifier. |
|  | data sources | 23 | Test for data sources that are no longer supported or replaced. |
|  | Wikidata | 15 | Test compatibility with Wikidata to support linked data. |
| COVID-19 |  | 2 | Tests that all interactions have a reference. |
| LIPID MAPS |  | 2 | Tests that only LIPID MAPS IDs are used for lipids. |
| IMD |  | 5 | Tests that enzyme identifiers are from UniProt, Ensembl, NCBI Gene, or Enzyme Nomenclature. |

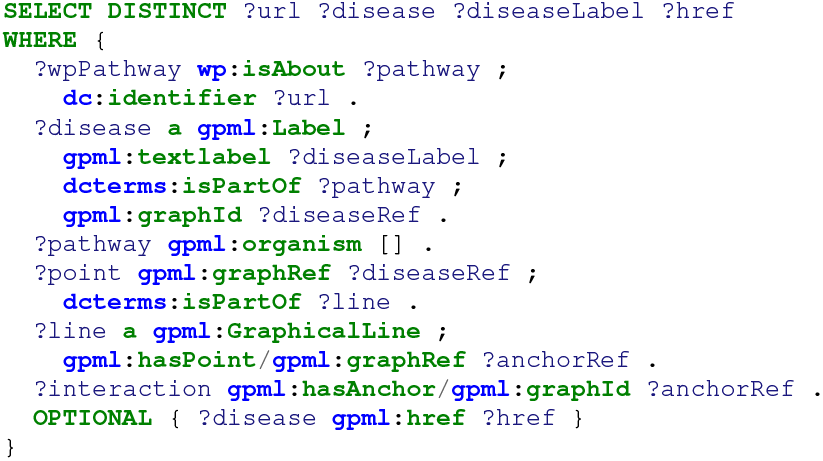

Code Example 1: Retrieve all disease labels in the IMD pathways and return the pathway URL (?url), disease (?disease and ?diseaseLabel), and link to OMIM (?href).

The curation tests were wrapped in JUnit (junit.org) test methods so that continuous integration platforms like Jenkins could easily run the tests and visualize the results. The SPARQL query codes could be run against the public WikiPathways SPARQL endpoint as well as on a local collection of GPML files.

Specifically for the IMD collection, several new tests were created, including non-numeric Rhea IDs; links to OMIM; if interaction IDs are from Rhea; if enzyme IDs are from UniProt, Ensembl, NCBI Gene, or Enzyme Nomenclature; and that all metabolites in the pathways are involved in at least one interaction.

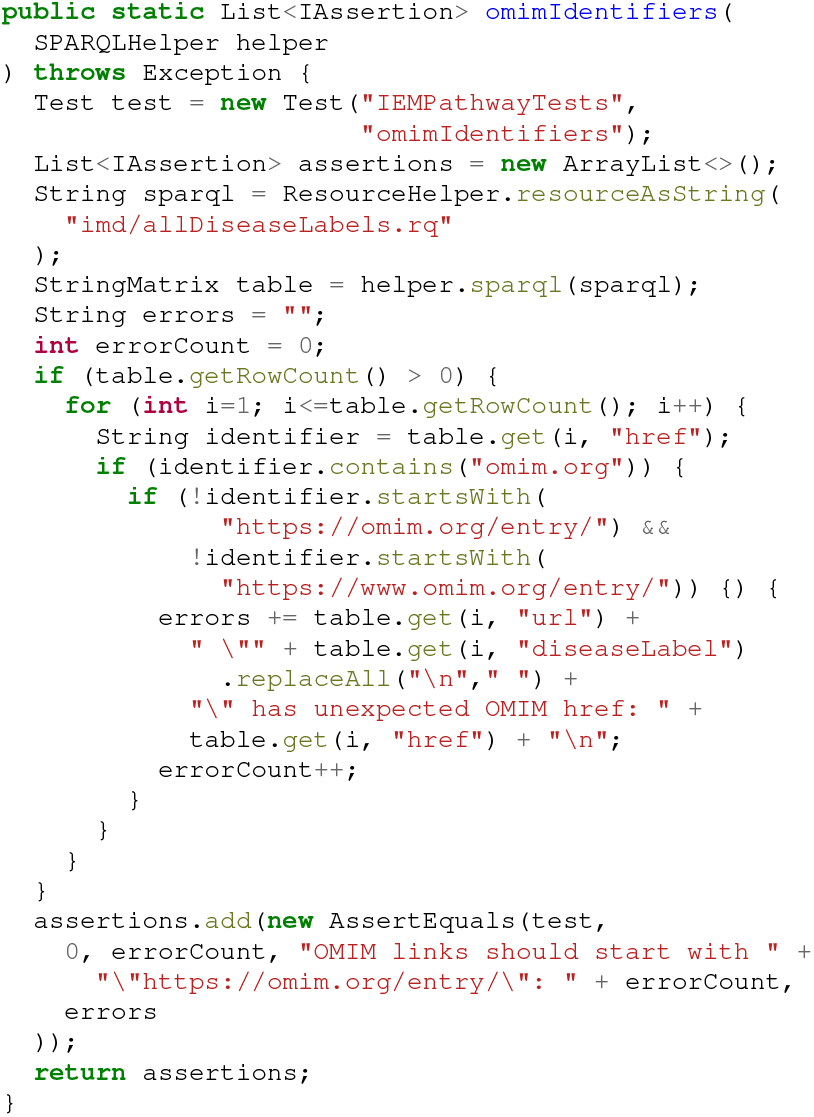

Code Example 2: Java code to analyze the results from a SPARQL query defined in the imd/allDiseaseLabels.rq SPARQL query; checks that links to OMIM have a resolving URL pattern (including the “/entry/” part). For each mismatch, an error is recorded and returned by the method.

### 1.2.6 A curation collection template

To support the development and curation of a specific collection of pathways, the RDF generation and data curation tools were combined to run a specific set of tests relevant to the collection. This approach has been shaped as a GitHub repository template (github.com/wikipathways/wikipathways-curation-template) with precompiled binaries of the GPML2RDF and WikiPathwaysCurator libraries, simplifying setting up data curation for custom pathway collections. Three Java utilities are provided, *CreateGPMLRDF, CreateRDF*, and *CheckRDF* to generate GPML-RDF, WP-RDF, and run the tests, respectively. The RDF generated is the same RDF model used for the monthly WikiPathways releases which are hosted in the SPARQL endpoint.

A *Makefile* automatically downloads newer versions of pathway models to allow running the tests regularly on the latest pathway models. The results of the curation tests were stored as Markdown files in the same GitHub repository. The template can be further configured by specifying on which website the Markdown files should be published in the *website*.*txt* file, and which curation tests should be run or not run with the *tests*.*txt* file.

### 1.2.7 Automated IMD pathway curation

Using this template (release 5, wikipathways-curation-template/releases/tag/release-5), a curation collection was set up for the IMD pathways at github.com/BiGCAT-UM/imd-pathway-curation. The collection was populated by listing the IMD pathways with the following SPARQL query:

The results were published using GitHub Pages at bigcat-um.github.io/imd-pathway-curation. This report also includes information on the Systems Biology Markup Language (SBML) format [40], which are obtained through a conversion process described elsewhere [41] through the MINERVA Conversion API [42].

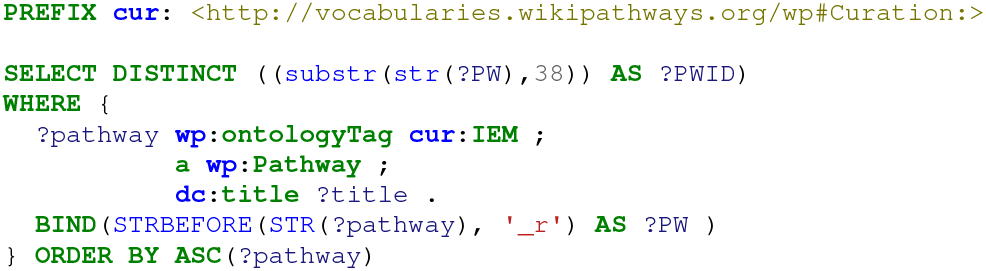

Code Example 3: Retrieve all WikiPathways IDs for IMD pathway models.

### 1.2.8 Querying the pathways

The content of the pathway models can be queried through the SPARQL-Virtuoso instance (sparql.wikipathways.org hosting the RDF data for pathway models with the quality tag “Approved for Data Analysis” [37]. For this study, data from April 2023 was used (archived at DOI:10.5281/zenodo.7853104). This section showcases several example queries to interact with the IMD pathway content through the SPARQL endpoint. These queries are available in GitHub (github.com/BiGCAT-UM/IMD Curation Analysis) and collect different information from the IMD pathway models. First, an overview of the data models and annotated DataNodes was obtained. Second, the IDs used in the IMD pathway models were investigated. Third, the content of the pathway models was compared to three pathway databases, KEGG [43], Reactome [44], and WikiPathways [30]. Fourth, the interaction types between DataNodes within the models were investigated. Last, the completeness of linked open data platforms such as Wikidata was checked, by performing a query to find IMDs based on the HGNC symbols of the proteins involved in the pathways (using the BridgeDb [25] for ID mapping in the WikiPathway RDF).

## 1.3 Results and Discussion

### 1.3.1 The GPML dataset

The complete list of 46 pathways can be found in the GitHub repository under github.com/BiGCAT-UM/imd-pathway-curation/blob/main/pathways.txt. An overview of the DataNode content in these IMD pathway models is depicted in Table 1.2. Many pathway models hold a combination of GeneProduct and protein DataNodes, which could cause complications downstream (e.g. uptake by Wikidata and ID harmonization). Each DataNode in a GPML model can be annotated with one database ID, which can be sourced from a plethora of databases (to name a few used to describe genes and proteins: HGNC symbol and number [45], Ensembl [46], NCBI (Entrez Gene) [47], Enzyme Nomenclature Code [48], InterPro [49], and UniProt [50]).

**Table 1.2:** Pathways and relevant content in the WikiPathways RDF. Uniqueness counts based on unification to Rhea IDs (metabolic reactions), and OMIM URLs (Diseases). Metabolic conversions which could not be connected to a Rhea ID are added in the respective column.

| Pathway title | PWID | Genes | Proteins | Metabolites | Rhea | No Rhea | Diseases |
| --- | --- | --- | --- | --- | --- | --- | --- |
| Degradation pathway of sphingolipids, including diseases | WP4153 | 14 | 21 | 18 | 5 | 10 | 19 |
| Biosynthesis and regeneration of tetrahydrobiopterin and catabolism of phenylalanine | WP4156 | 4 | 21 | 32 | 15 | 30 | 11 |
| GABA metabolism (aka GHB) | WP4157 | 5 | 20 | 28 | 18 | 19 | 2 |
| Neurotransmitter disorders | WP4220 | 8 | 11 | 23 | 7 | 16 | 6 |
| Purine metabolism and related disorders | WP4224 | 2 | 25 | 64 | 58 | 8 | 15 |
| Pyrimidine metabolism and related diseases | WP4225 | 5 | 29 | 44 | 47 | 11 | 11 |
| Vitamin B6-dependent and responsive disorders | WP4228 | 3 | 7 | 20 | 4 | 16 | 5 |
| Krebs cycle disorders | WP4236 | 7 | 18 | 23 | 16 | 24 | 4 |
| Disorders of folate metabolism and transport | WP4259 | 1 | 20 | 30 | 8 | 19 | 12 |
| Vitamin B12 disorders | WP4271 | 2 | 16 | 19 | 2 | 9 | 12 |
| MTHFR deficiency | WP4288 | 11 | 19 | 23 | 7 | 11 | 1 |
| Methionine metabolism leading to sulfur amino acids and related disorders | WP4292 | 6 | 12 | 23 | 6 | 12 | 7 |
| Thiamine metabolic pathways | WP4297 | 8 | 1 | 22 | 0 | 15 | 3 |
| Cysteine and methionine catabolism | WP4504 | 5 | 16 | 39 | 16 | 15 | 0 |
| Tyrosine metabolism and related disorders | WP4506 | 1 | 9 | 24 | 8 | 12 | 6 |
| Molybdenum cofactor (Moco) biosynthesis | WP4507 | 1 | 8 | 11 | 1 | 5 | 5 |
| Gamma-glutamyl cycle for the biosynthesis and degradation of glutathione, including diseases | WP4518 | 2 | 6 | 11 | 7 | 0 | 5 |
| Cerebral organic acidurias, including diseases | WP4519 | 2 | 8 | 28 | 10 | 8 | 5 |
| Glycosylation and related congenital defects | WP4521 | 1 | 25 | 29 | 18 | 2 | 21 |
| Metabolic pathway of LDL, HDL and TG, including diseases | WP4522 | 10 | 17 | 5 | 0 | 0 | 17 |
| Classical pathway of steroidogenesis with glucocorticoid and mineralocorticoid metabolism | WP4523 | 13 | 22 | 30 | 29 | 15 | 14 |
| Alternative pathway of fetal androgen synthesis | WP4524 | 12 | 13 | 18 | 14 | 9 | 7 |
| Oxysterols derived from cholesterol | WP4545 | 19 | 40 | 62 | 8 | 61 | 12 |
| Urea cycle and associated pathways | WP4595 | 2 | 24 | 30 | 23 | 4 | 11 |
| Leucine, isoleucine and valine metabolism | WP4686 | 3 | 25 | 74 | 23 | 3 | 21 |
| Serine metabolism | WP4688 | 1 | 8 | 26 | 7 | 6 | 0 |
| Pathways of nucleic acid metabolism and innate immune sensing | WP4705 | 7 | 19 | 4 | 2 | 0 | 0 |
| Sphingolipid metabolism overview | WP4725 | 20 | 23 | 26 | 4 | 23 | 1 |
| Purine metabolism | WP4792 | 2 | 15 | 37 | 47 | 6 | 0 |
| Phosphoinositides metabolism | WP4971 | 4 | 54 | 15 | 24 | 4 | 4 |
| Amino acid transport defects (IEMs) | WP5029 | 0 | 9 | 27 | 0 | 0 | 5 |
| Ethylmalonic encephalopathy | WP5030 | 1 | 4 | 12 | 3 | 6 | 1 |
| Biotin metabolism, including IMDs | WP5031 | 1 | 9 | 15 | 9 | 9 | 5 |
| Riboflavin and CoQ disorders | WP5037 | 2 | 16 | 27 | 6 | 8 | 10 |
| 7-oxo-C and 7-beta-HC pathways | WP5064 | 14 | 17 | 27 | 1 | 40 | 4 |
| Glyoxylate metabolism | WP5166 | 1 | 14 | 16 | 9 | 2 | 3 |
| Hemesynthesis defects and porphyrias | WP5169 | 9 | 0 | 18 | 2 | 8 | 10 |
| Leukotriene metabolic pathway | WP5171 | 13 | 4 | 19 | 11 | 7 | 5 |
| Disorders of galactose metabolism | WP5173 | 13 | 3 | 10 | 3 | 3 | 13 |
| Disorders in ketone body synthesis | WP5175 | 0 | 5 | 13 | 1 | 2 | 4 |
| Disorders of bile acid synthesis and biliary transport including diseases | WP5176 | 18 | 6 | 44 | 4 | 7 | 17 |
| Disorders of fructose metabolism | WP5178 | 14 | 0 | 17 | 3 | 1 | 14 |
| Copper metabolism | WP5189 | 4 | 0 | 2 | 0 | 0 | 5 |
| Creatine pathway | WP5190 | 4 | 2 | 18 | 5 | 2 | 6 |
| Cholesterol synthesis disorders | WP5193 | 18 | 12 | 17 | 3 | 3 | 18 |
| Disorders in ketolysis | WP5195 | 0 | 5 | 8 | 1 | 2 | 3 |
| Ether lipid biosynthesis | WP5275 | 3 | 22 | 17 | 10 | 4 | 7 |

Various IDs are used within the pathway models, see Table 1.3. These results show that Ensembl is used for most annotations regarding GeneProduct DataNodes, UniProt for Proteins, and ChEBI for Metabolites. Furthermore, several pathways contain cross-references to other pathways (65 in total). The highlighted section in this Table shows a misannotation (a DataNode with type “Metabolite”, however with annotation from Entrez Gene). The pathway containing a wrong annotation can be easily retrieved (WP4220)and curated (which we performed on the 20th of April 2023).

**Table 1.3:** Identifiers (IDs) used in IMD pathway models, split up by DataNode type, providing the database source, number of identifiers from that source, and the type of biological entity, respectively. The row highlighted in yellow indicates an annotation that needs reviewing by a curator.

| datasource | numberEntries | dataNodes |
| --- | --- | --- |
| Wikidata | 8 | Rna |
| Uniprot-TrEMBL | 430 | Protein |
| Enzyme Nomenclature | 59 | Protein |
| Ensembl | 49 | Protein |
| InterPro | 6 | Protein |
| Entrez Gene | 6 | Protein |
| Reactome | 3 | Protein |
| KEGG Genes | 1 | Protein |
| WikiPathways | 65 | Pathway |
| ChEBI | 639 | Metabolite |
| LIPID MAPS | 85 | Metabolite |
| Wikidata | 58 | Metabolite |
| HMDB | 39 | Metabolite |
| InChIKey | 30 | Metabolite |
| PubChem-compound | 26 | Metabolite |
| CAS | 8 | Metabolite |
| Chemspider | 1 | Metabolite |
| Entrez Gene | 1 | Metabolite |
| Ensembl | 117 | GeneProduct |
| HGNC | 20 | GeneProduct |
| Entrez Gene | 19 | GeneProduct |
| Uniprot-TrEMBL | 10 | GeneProduct |
| BRENDA | 2 | GeneProduct |
| Enzyme Nomenclature | 2 | GeneProduct |
| InterPro | 1 | GeneProduct |

Comparing the data captured with the IMD models revealed that novel content was added for both metabolites and gene content. Figure 1.3 shows the overlapping content between human pathway models from KEGG, Reactome, WikiPathways, and the IMD pathway models. The content in the IMD models is much smaller compared to all pathway model content in these three databases, however, several novel metabolites and genes were added through these pathways. These results highlight the importance of capturing IMD pathways in computational models and the unique data captured through these models. Comparing the different interactions between metabolites is not directly possible at this moment, due to differences in annotations between the directionality of the interactions by the aforementioned databases.

**Figure 1.3:**
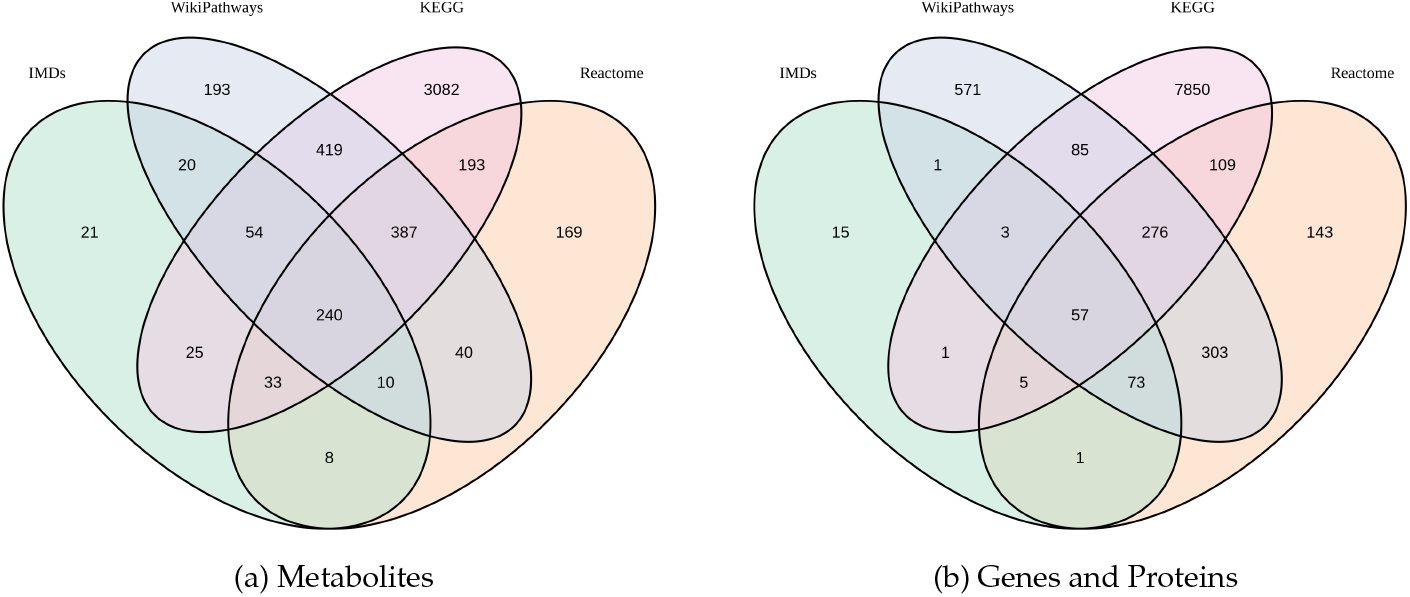
Overview of metabolite and gene/protein pathway content from three pathway databases (KEGG, Reactome, WikiPathways) compared to the content from the IMD models. Metabolites were unified to the KEGG Compound ID, genes and proteins to the Entrez (NCBI) gene ID using BridgeDb.

A similar query was performed on annotations for interactions, to understand which types of DataNodes are used within the pathway models on IMDs. The results of this query are depicted in Table 1.4. As expected, most of the interactions are metabolic conversion between two Metabolite DataNodes. The other interactions contain a variety of Stimulation, Transcription-Translation, and Inhibitions. Enzymes catalyzing a reaction are connected to an anchor on a metabolic conversion in the PathVisio pathway models, therefore this query cannot reflect the number of metabolic conversion reactions including an enzyme catalyzing the reaction.

**Table 1.4:** Interaction types used in the WikiPathways IMD models, including their source and target DataNode type. Column names follow matching variables in the SPARQL queries and count the MIM-interaction types for each unique combination of source and target types.

| Count-Interactions | MIMTypes | sourceType | targetType |
| --- | --- | --- | --- |
| 812 | Conversion | Metabolite | Metabolite |
| 13 | Stimulation | Protein | Protein |
| 7 | Stimulation | Protein | GeneProduct |
| 6 | TranscriptionTranslation | Rna | Rna |
| 6 | Stimulation | Metabolite | Protein |
| 4 | TranscriptionTranslation | GeneProduct | Protein |
| 4 | Stimulation | Rna | Protein |
| 3 | TranscriptionTranslation | GeneProduct | Rna |
| 3 | Inhibition | Metabolite | GeneProduct |
| 3 | Inhibition | Metabolite | Protein |
| 2 | Inhibition | Protein | Rna |
| 2 | Stimulation | GeneProduct | Protein |
| 2 | TranscriptionTranslation | Protein | Protein |
| 2 | Stimulation | GeneProduct | GeneProduct |
| 2 | Inhibition | Metabolite | Metabolite |
| 2 | Stimulation | Rna | GeneProduct |
| 1 | TranscriptionTranslation | GeneProduct | GeneProduct |
| 1 | Conversion | Rna | Rna |
| 1 | Inhibition | Protein | GeneProduct |
| 1 | Stimulation | Metabolite | GeneProduct |

In order to evaluate the completeness of linked open data platforms a federated query was performed against Wikidata to find IMDs based on the HGNC symbols of the proteins involved in the pathways. Out of the 32 queried proteins, eight returned a result regarding a connection to a disorder in Wikidata listed as an IMD (Table 1.5). Again these results show the importance of capturing data on rare metabolic disorders through pathway models since these models can be used to collect information on a process level rather than on an individual level. By capturing this data in a structure that can be used for Linked Open Data approaches, other researchers can easily evaluate which information is present (and also lacking!). Furthermore, using the community curation platform WikiPathways allows users to add missing data or update existing models with novel discoveries.

**Table 1.5:**
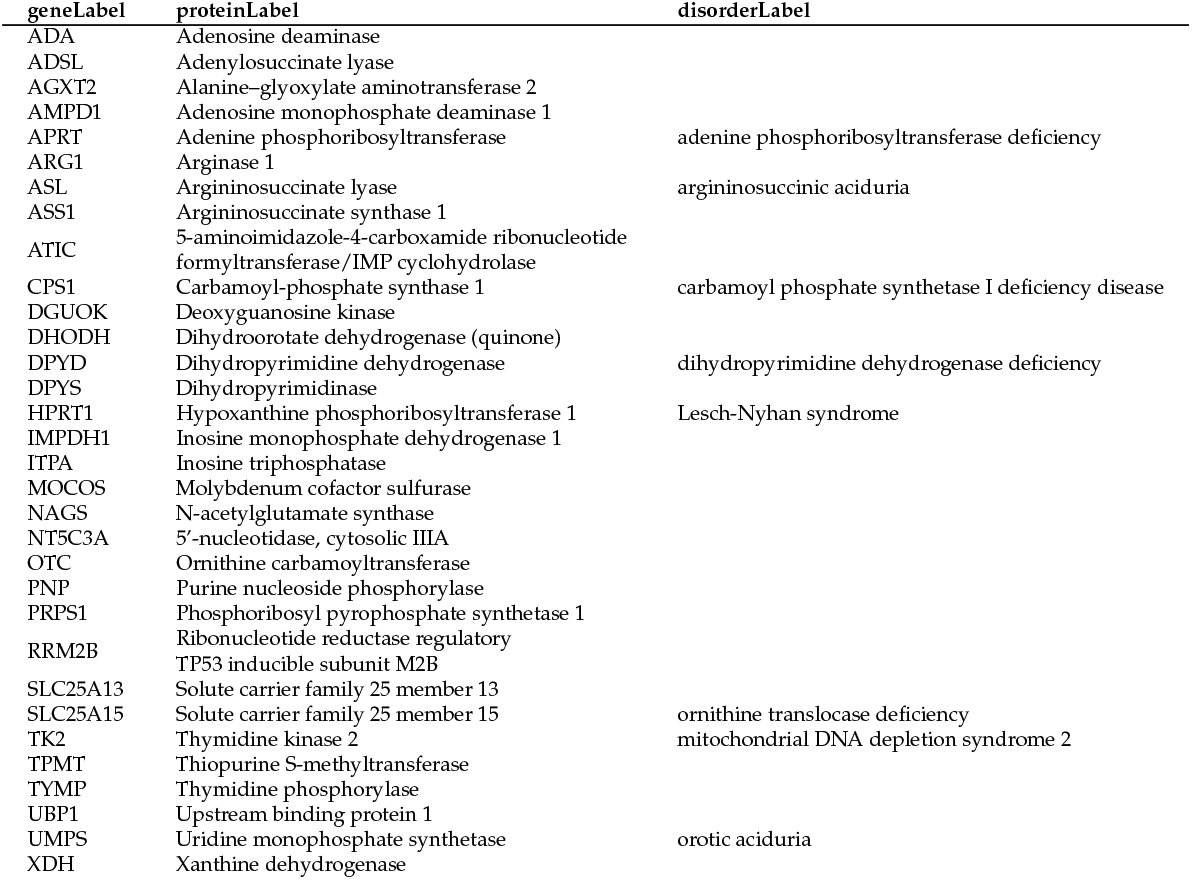
An overview of Genes from the Purine (WP4224), Pyridimide (WP4225), and Urea Cycle (WP4595) IMD pathway models. Column names follow matching variables in the SPARQL queries and provide IMD data captured in Wikidata.

### 1.3.2 Limitations

Several pathway models contain IDs describing an enzyme family (InterPro, Enzyme Nomenclature), which can create issues mapping data to these individual DataNodes. However, often the general understanding of the underlying biology of these proteins involved in IMDs is lacking and therefore these entries cannot be modeled with more precision. The knowledge currently captured in so-called ‘groups’ (e.g. two enzymes catalyzing the same chemical conversion, without forming a complex) is currently not taken up in the RDF model and therefore not queryable.

Currently, the disease annotations are based on URLs from OMIM and not added to the models as DataNode annotations. The information on the disease nodes is part of the RDF for querying, but not in a unified and harmonized format. the current PathVisio GPML model is not directly suitable for disease annotations, however, this functionality will be added in a future version. Support for rich disease annotation on a DataNode level, for example adding the Human Disease Ontology [51] or the Evidence and Conclusion Ontology [52], could allow for a more precise annotation of curation decisions.

## 1.4 Conclusions

The community curation of Inherited Metabolic Disorders (IMDs) pathways has led to the development of 47 machine-readable pathway models supporting Linked Open Data approaches and the description of several novel metabolic interactions. Additional curation tests were developed to evaluate the content within these models using a combination of SPARQL queries and JUnit tests. The results from this project were integrated into updates in the WikiPathways RDF, as well as support disease DataNode types and annotations in future versions of PathVisio.

